# Potential nematicides identified through targeted *in vivo* screening of a 5-HT-gated chloride channel, MOD-1

**DOI:** 10.64898/2026.01.25.701312

**Authors:** Fernando Calahorro, Yogendra K. Gaihre, Katarzyna Marek, Claude Charvet, Cedric Neveu, Mirela Coke, Catherine Lilley, Peter Urwin, Lindy Holden-Dye, Vincent O’Connor

**Affiliations:** School of Biological Sciences, Life Sciences Building 85, University of Southampton (UK); INRAE, Université de Tours (France); Centre for Plant Sciences, School of Biology, University of Leeds (UK); MSD Animal Health Innovation GmbH (Germany)

**Keywords:** *Globodera pallida*, *Caenorhabditis elegans*, plant parasitic nematode, pharynx, *Xenopus* oocyte, anthelmintic, nematicide

## Abstract

New approaches to mitigate the reduction of crop yields by plant parasitic nematodes are needed in the face of increasing concerns of the impact of nematicides on precious ecosystems. One approach is to target receptors in the parasitic nematode that are vital for their survival that less widely expressed in non-target organisms. Nematodes express a phylogenetically restricted 5-HT-gated chloride channel, MOD-1, activation of which causes paralysis in *Caenorhabditis elegans.* We show that MOD-1 is expressed in the motor nervous system of the plant parasitic nematode *Globodera pallida* and its functional characterisation is validated by 5-HT activation when reconstituted in *Xenopus laevis* oocytes. To evaluate MOD-1 as a nematicide target we utilised a previously described platform called ‘PhaGeM4’ for ‘<u>Pha</u>rmaco<u>Ge</u>netic targeting of <u>M4</u> neurone’ in which MOD-1 is expressed in transgenic *C. elegans* and nematode development in the face of MOD-1 chemical modulation is tracked. We screened Pathogen Box, a chemical library of 400 diverse drug-like molecules, using PhaGeM4. This identified 10 putative ‘hits’ for *C. elegans* MOD-1. These hits were pursued through a sequential, iterative pipeline encompassing *mod-1* dependent *C. elegans* motility and G.*pallida* motility assays in combination with pharmacological interrogation of *G. pallida mod-1* in PhaGeM4. This approach highlights 3 compounds with a *mod-1* dependent action (quipazine, our benchmark compound; MMV687251, a vancomycin-like compound; MMV688774, an antifungal with common name posaconazole) and one compound that acts through an undetermined target (MMV002816, also known as the antifilarial drug, diethylcarbamazine). Each of these compounds had a significant inhibitory effect on *G. pallida* J2 root invasion. Overall, this lends confidence that the PhaGeM4 screening platform can delivery new chemical leads for crop protection and highlights four new chemistries of interest. More generally, this approach could be applied to other ligand-gated ion channels of interest as targets.

## INTRODUCTION

Plant parasitic nematodes cause major crop losses and for more than a century chemical nematicides have played a significant role in limiting their impact [1]. However, increasing concerns regarding the eco-toxicity of the most widely used and successful nematicides, and resulting regulatory controls limiting or banning their application has energised renewed efforts to develop alternatives. In terms of nematicides, this encompasses new chemistries that have reduced toxicity to non-target organisms that have seen the development of fluopyram [2], fluazaindolazine [3] and fluensulfone [4]. For fluensulfone, uniquely, this selectivity also appears to spare free-living nematodes, as *Caenorhabditis elegans* has a low sensitivity compared to *Meloidogyne sp.* [5] highlighting the potential for phyla selective mode of actions.

Further efforts to identify economically viable, ‘new generation’ nematicides are required to address the significant challenge posed by plant parasitic nematodes to food security. Recently, the potential of biogenic amine signalling as an attractive target for highly selective chemical control of parasitic nematodes has been proposed [6]. 5-HT (5-hydroxytryptamine, 5-HT), paralyses *C. elegans* and this effect is dependent on *mod-1*, a 5-HT-gated chloride channel that is expressed in the motor nervous system [7]. The MOD-1 receptor shows an unusual phylogenetically constrained distribution and appears to be largely restricted to the phylum Nematoda. It is not found in insects e.g. in bees, and not at all in mammals [8]. This highlights that chemicals that act selectively on the MOD-1 receptor in plant parasitic nematodes would have an excellent potential for selective toxicity. Recent indications of the pharmacological profile of MOD-1, also support this receptor as a potential target for animal parasitic nematodes [8, 9].

We previously developed a screening platform that could be used to identify chemical modulators of invertebrate ion channels including MOD-1 [8]. In this platform, we made optimum use of the experimental tractability of *C. elegans*, to transgenically express *mod-1*, cloned from *C. elegans* in a single pharyngeal neurone, M4. Hence the name, PhaGeM4, for ‘Pharmacogenetic targeting of M4 neurone’. This neurone was chosen as earlier studies have shown that if this neurone is subject to laser ablation during *C. elegans* development the worms undergo developmental arrest and die [10]. This approach was benchmarked using *mod-1* and we showed that application of 5-HT to transgenic *C. elegans* expressing *mod-1* in M4 phenocopied the laser ablation causing L1 arrest. This work thus delivered an an assay for chemicals that modulate MOD-1 [8].

In the present work, we utilise PhaGeM4 with respect to MOD-1 for identifying selective compounds. Importantly, we show that *mod-1* is expressed in the motor nervous system of a major pest, the potato cyst nematode *G. pallida* with a similar pattern to that observed in *C. elegans* [7] suggesting it may also underpin similar functional roles. We confirm this as a bona fide 5-HT-gated chloride channel through expression in *Xenopus laevis* and then exploit the PhaGeM4 platform to resolve chemicals from compound libraries that modulate MOD-1. For this we used the Pathogen Box library (https://www.pathogenbox.org/) which consists of 400 drug-like molecules active against neglected diseases of interest such as cryptosporidiosis, trichuriasis, tuberculosis, kinetoplastids, malaria, schistosomiasis, filariasis and Dengue virus [11]. This library has previously been exploited as a powerful tool for repurposing compounds in high-throughput screening approaches [12]. The results of our pipeline support the contention that screening compound libraries using the PhaGeM4 platform has potential to identify chemical modulators of MOD-1 that impact root invasion of *G. pallida* J2s and has delivered at least three new chemistries of interest.

## METHODS

### MOD-1 protein topology prediction

The distinct protein sequences that are predicted to encode *C. elegans* and *G. pallida* MOD-1 were subject to transmembrane topology analysis using Protter (version 1.0) with default settings (Omasits et al, 2014).

### *In situ* hybridization

A set of 30 HCR probes designed to the coding region (Molecular Instruments) and amplifiers labelled with Alexa 647 were used to detect the expression pattern of *G. pallida mod-1*. *G. pallida* J2s were fixed, sectioned and permeabilised as described previously [13]. The subsequent *in situ* hybridization followed protocols described previously [14, 15]. The permeabilised J2 sections were washed for 5 min in 1:1 Hybridisation Buffer (Molecular Instruments) and 1% PBST (PBS with 1% Tween). The nematodes were pre-hybridized in Hybridization Buffer (Molecular Instruments) for 30 min prior to incubation overnight in hybridization buffer with 1% probe stock solution. The J2s were washed three times, 15 min each time, in 37°C pre-warmed Probe Wash Buffer (Molecular Instruments) and once for 5 min in 5xSSCT with 0.1% Tween prior to incubation for 30 min in amplification buffer (Molecular Instruments). The nematodes were then incubated for 20-22 hours at room temperature with the amplification hairpins prepared as detailed in Choi et al (2016). The nematodes were washed prior to re-suspension in mounting medium (Slow Fade Gold antifade mountant). Confocal microscopy (LSM 800 confocal microscope connected to ZEN software) was used to capture maximum fluorescence projection images which were superimposed on a brightfield image.

### Drugs and chemicals

#### MMV Pathogen Box: Format and storage

The original collection of 400 compounds within MMV Pathogen Box V2 was supplied in dissolved in 100 % DMSO in four sealed 96-wells microtiter plates (A-E, 80 compounds in each plate at a concentration of 10 mM in a volume of 10 µl. The library shipped on dry ice and stored at −80⁰C until use. For *post hoc* validation of primary experiments and re-testing of ‘hits’ ∼ 2mg lyophilized samples were received from Evotec Toulouse (France) and resuspended in DMSO (100%) as 50 mM stocks and stored at −80⁰C until use.

### *C. elegans* strains and culture conditions

The culture and maintenance of *C. elegans* were as previously described [16]. Monoxenic cultures of *C. elegans* embryos in S Medium based liquid cultures were as described [8].

Bristol N2 wild type (wt) and MT9668 *mod-1* (*ok103*) *V* x6 outcrossed strains were provided by CGC (*Caenorhabditis* Genetics Center, University of Minnesota). For the Pathogen Box screening performed in this study, the following worm strains were used as described previously [8]:

N2 (wt) sIs [P*ceh-28*::Ce*mod-1;* P*myo-3::gfp*] x3 outcrossed (*mod-1 C. elegans* version)
N2 (wt) sIs [P*ceh-28*::Gp *mod-1;* P*myo-3::gfp*] x3 outcrossed (*mod-1 G. pallida* version)

P*ceh-28* is a promoter sequence that drives selective expression of the transgene in the *C. elegans* pharyngeal neurone M4.

### *Globodera pallida* maintenance and culture

Cysts of *G. pallida* population Lindley (Pa2/3) were extracted from infested sand/loam pre -cultivated with potato plants *cv*. Desiree, using the floatation technique. The cysts were sterilized in 0.1% chlorhexidine digluconate, 0.5 mg/ml CTAB for 30 mins and washed once with filter sterilized tap water, before being exposed to potato root diffusate at 20°C to stimulate hatching, for the nematode invasion assays. For all the other assays, dry cysts were treated with 0.1% malachite green solution for 30 mins for an initial sterilisation step, followed by extensive washing in tap water. Cysts were then incubated in an antibiotic cocktail [17] at 4°C overnight and washed five times with sterile tap water. To induce hatching, cysts were placed in a solution of 1 part potato root diffusate to 3 parts tap water. The root diffusate was obtained by soaking washed roots of three-week-old potato plants in tap water at room temperature for 3 hours, using a ratio of 80 g root/litre. The diffusate was then filter-sterilised and stored at 4°C. *G. pallida* J2s will typically begin hatching one week after exposure to potato root diffusate. *G. pallida* J2s collected over one week were used in root invasion assays, while J2s that had hatched within the previous 24 h were used for all the other experiments. Prior to the experiments the J2s were washed in pure water to remove the potato root diffusate.

### *Xenopus laevis* oocyte electrophysiology

To functionally characterise MOD-1 from both *C. elegans* and *G. pallida*, cRNAs were transcribed *in vitro* from plasmid containing the inserts encoding the cDNA for *C. elegans* and *G. pallida mod-1* using the mMessage mMachine T7 transcription kit (Thermofisher). Receptors were reconstituted in *Xenopus laevis* oocytes and assayed under voltage-clamp as previously described [18]. Briefly, 36 nl of cRNA mix were microinjected in defolliculated *Xenopu*s oocytes (Ecocyte Bioscience) using a Nanoject II microinjector (Drummond). After 4 days incubation, oocytes were voltage-clamped at a holding potential of −70 mV and electrophysiological recordings were carried out as described previously [18]. Whole cell current responses were collected and analysed using the pCLAMP 10.4 package (Molecular Devices).

### Harvesting *C. elegans* embryos

Embryos were isolated as previously described [8]. Briefly, approximately 600 gravid adults were washed with sterile ddH_2_O. The washed worms were pelleted and lysed using 12% alkaline hypochlorite solution in 1.6 M NaOH. The lysis was halted with egg buffer (final concentration of 25 mM HEPES, 118 mM NaCl, 48 mM KCl, 2 mM CaCl_2_, 2 mM MgCl_2_) and the embryos pelleted. Then the egg pellet was resuspended by mild agitation in 2.5 ml of ddH_2_O. 2.5 ml sterile 60% sucrose (W/V) was carefully added below the eggs and centrifuged for 1 min at 1200 rpm. Approximately 3 ml of the sucrose harbouring the embryos was transferred into a fresh sterile 15 ml conical tube and washed with 5 ml of ddH_2_O and centrifuged for 6 min at 1200 rpm. After two further washes the isolated egg pellet was resuspended in 100 µl of sterile ddH_2_O. Finally, three 1 µl aliquots were transferred onto agar plates and the number of eggs counted using a binocular dissecting microscope (Nikon SMZ800) to obtain an indication of the concentration of eggs in the solution.

### Behavioural assays

#### Testing Pathogen Box library in the PhaGeM4 C. elegans developmental assay

The *C*. *elegans* developmental assay that underpins the PhaGeM4 bioassay were performed as previously described [8]. Briefly, ∼20 eggs from the indicated transgenic strain were dispensed into individual wells of 96-well microtiter plates containing 50 µl of monoxenic culture. The number of eggs dispensed into each well was recorded. A volume of 0.125 µl (0.375 µl in 150 µl of monoxenic culture to aliquot three replicas) stocks of each compound in Pathogen Box library, at its original concentration (10 mM, 100% DMSO), was added to the specified volume of monoxenic culture to a final concentration of 25 µM (0.25% DMSO). The eggs developed in the presence of the indicated concentration of compounds, and fluorescence driven by the co-expressed body wall muscle GFP was monitored from each well at 72 hrs using a FLUOstar microplate reader (BMG Labtech) providing a proxy for the developmental progression of the embryos through to adult.

To assess the transgene dependent effect for each compound, we tested in parallel the development of both *Pceh-28::mod-1* transgenic and N2 (wt) expressing *myo-3 gfp* animals. The effect on the N2 strain was assessed by visual inspection after 72 hrs of incubation in each compound. Compounds were identified as having no detrimental effect on N2 development if the embryos developed beyond the L4 developmental stage to adults after 72 hrs.

#### Further characterisation of PhaGeM4 “hits” in a C. elegans thrashing assay

In liquid *C. elegans* exhibits a stereotyped swimming motion called ‘thrashing’. We conducted population-based thrashing assays that assess the pharmacological effect of compounds on motility. 10 to 20 *C. elegans* (L4+1 day old), grown on OP50 NGM plates at 20 ⁰C, were picked into individual wells of a sterile Corning® 96-well microplate (Clear Flat Bottom Polystyrene), containing 200 μl M9 buffer with either vehicle or test compound. These experiments were conducted on N2 (wt) *C. elegans* and *mod-1(ok103)*. The number of immotile worms in each well was scored after the indicated time of exposure and expressed as a percentage of the total number of worms in the well. An animal was considered immotile if it did not exhibit any thrashing motion for a period of 5 s.

#### Further characterisation of PhaGeM4 “hits” in a G. pallida motility assay

*G. pallida* cysts set up in potato root diffusate (PRD) in 30 mm Petri dishes from 1 to 3 weeks were used to obtain J2s. The J2s that emerge from the cyst overnight in fresh PRD were transferred into 30 mm Petri dishes and washed with ddH_2_O (0.01 % BSA) for 10 min. Then ∼ 10 J2s were transferred into individual wells of a sterile Corning® 96-well microplate (Clear Flat Bottom Polystyrene) containing either 50 µM of the compound ‘hits’ or vehicle (0.1% DMSO) in 200 µl of water. Individual worms were observed for 10 s, after 4 hrs and 24 hrs of incubation. J2s were defined as immotile if they held a straight body posture and were rod shaped within the 10 s observation period.

#### Further characterisation of PhaGeM4 “hits” in a G. pallida root invasion assay

*G. pallida* J2s obtained as described above, were collected every second days, and kept at 11 °C. The day prior to the inoculation of the plant root culture they were treated with the ‘hit’ compounds to be tested, or, in case of control J2s with water, 0.1% DMSO, 0.3% DMSO or 0.9% DMSO, as appropriate. For these assays the lyophilized test compounds were dissolved in 100% DMSO. As the compounds display differential solubilities the following maximal stock concentrations were achieved and are indicated. B25 (450 µM), B52 (50 µM), B59 (50 µM), B78 (150 µM), C25 (150 µM), E20 (50 µM), E26 (50 µM), E34 (150 µM), E42 (450 µM), E58 (150 µM). These stocks were stored at −20°C prior to use. Quipazine was dissolved in distilled water each time prior to use.

For root invasion assays, 6-8 rooted potato plants per treatment or control were transferred from *in vitro* culture to growth pouches (Mega International) and maintained at 20 °C with a 16:8 hr light:dark cycle. After 7 days, five root tips per root system were inoculated with ∼ 25 *G. pallida* J2s/root tip. A piece of Gf/a paper soaked in the appropriate compound solution was placed at the inoculation point. At 7 days post infection, the roots were stained with acid fuchsin to visualize and count total number of nematodes for each plant. The invasion assays for each compound were carried out on at least two separate occasions.

#### Data analysis

Bar graphs are presented as the mean ± standard error of the mean, with individual data points represented as scatter graphs, for the number of experiments as shown in individual figures. For concentration-response curves, data points were plotted as the mean ± standard error of the mean for the number of experiments shown in individual figures. Data were plotted using GraphPad Prism 7.01 software (San Diego, California). Statistical significance was determined either by unpaired Student’s t-test, one-way or two-way ANOVA as appropriate; significance level set at P< 0.05, followed by Bonferroni multiple comparisons as appropriate. For the oocyte experiments, responses were normalised to the response to 1 µM 5-HT. EC_50_ values with 95% confidence intervals were determined using GraphPad Prism 7.01 by plotting log concentration agonist against response and fitting the data to the equation; Y=Bottom + (Top-Bottom)/(1+10^((LogEC50-X)). For analysis of the fluorescence measurements in the PhaGeM4 *C. elegans* developmental assay we calculated a fluorescence intensity relative to control as follows: After scanning the plate, we obtained a signal value for each well. This value was divided by the number of eggs initially placed in the well to calculate fluorescence per egg. To normalise this to control, we divided the fluorescence per egg value for the test compound by the fluorescence per egg value for the control (non-drug).

#### Other reagents

5-HT hydrochloride (5-HT) (Merck-SigmaAldrich; CAS number: 153–98-0); Quipazine (QPZ) (Merck-SigmaAldrich; CAS number: 5786-68-5); Piperazine citrate (Merck-SigmaAldrich; CAS number 41372-10-5). For these additional compounds stocks used in the pharmacological experiments were made from powder, dissolved in water, protected from light and added to the experiment on the day of use.

## RESULTS

### MOD-1 topology and expression pattern in *G. pallida*

The topology of MOD-1 for both *C. elegans* and *G. pallida* amino acids reveal four transmembrane domains with an intracellular motif between transmembrane domains 3 and 4. This intracellular domain is larger in MOD-1 *G. pallida* (Figure 1). To rationalise the aim of targeting MOD-1 to mitigate root invasion by plant parasitic nematodes, we determined the expression pattern of *G. pallida mod-1* (*Gpa-mod-1*) in J2 nematodes using fluorescent *in situ* hybridisation. This revealed expression in several cell bodies in the head (Figure 1A,B), the ventral cord (Figure 1C) and the tail of *G. pallida J2s* (Figure 1D), a pattern similar to that described for adult hermaphrodite *C. elegans mod-1* [19]. The expression of *mod-1* in cell bodies around the metacorpus and along the ventral cord likely underpin its involvement in multiple behaviours critical to the plant parasitic nematode life cycle including motility, stylet thrusting and pharyngeal pumping.

**Figure 1.**
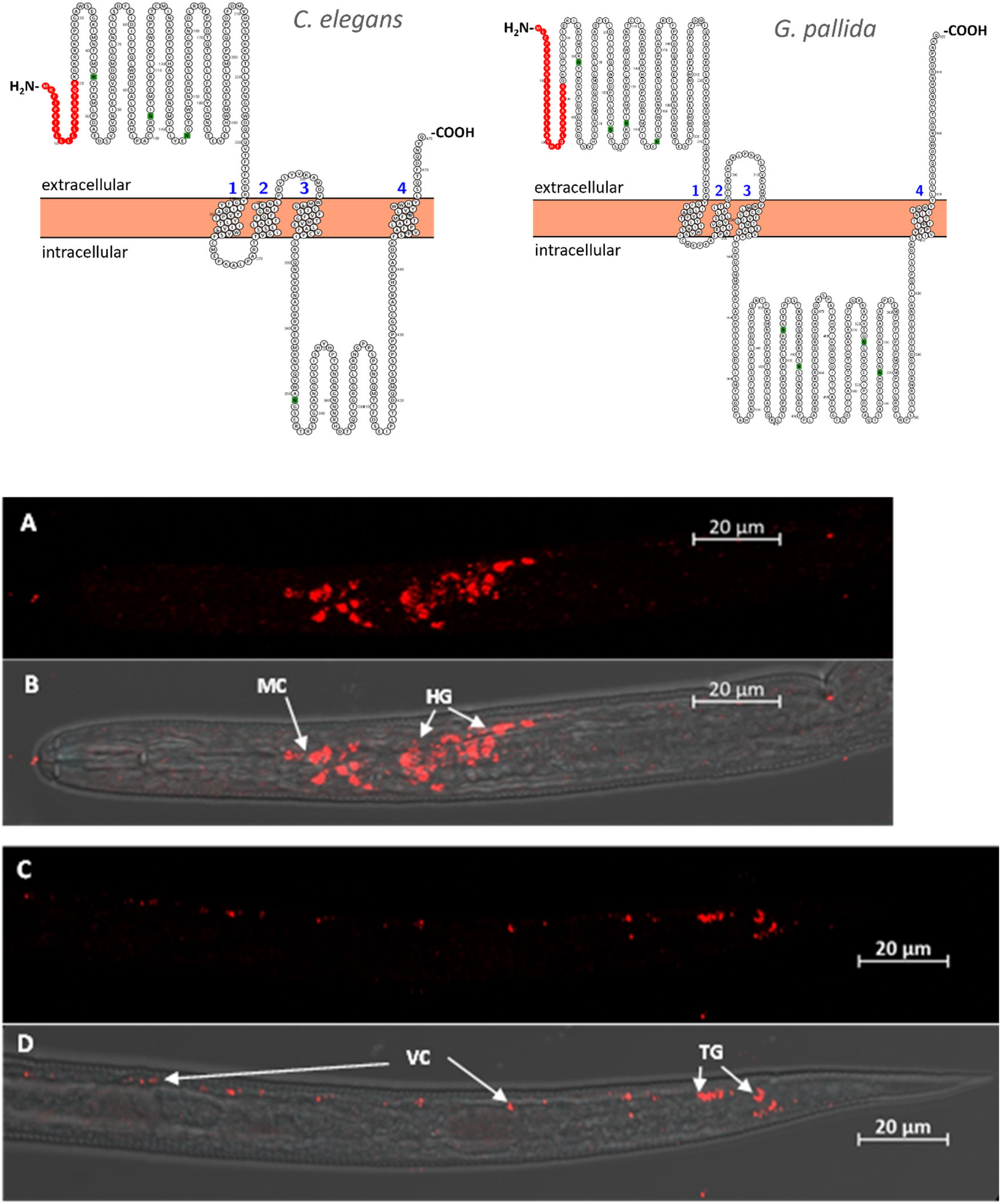
MOD-1 is expressed in *G. pallida*. **Top panel:** Predicted topology for versions from *C. elegans* and *G. pallida* MOD-1: MOD-1 *C. elegans* and *G. pallida* amino acids topology organization reveals four transmembrane domains with an intracellular motif between transmembrane domains 3 and 4. Interestingly, this intracellular domain is larger in MOD-1 *G. pallida*. Numbers indicate transmembrane domains. Residues in red indicate signal peptide and residues in green indicate N-glycosylated motifs. **Bottom panel:** Localisation of *G. pallida mod-1* transcripts. **A, B** The expression of *mod-1* was localized in a number of cell bodies around the metacorpus (MC) area and in the head ganglia (HG), but also **C, D** along the ventral cord (VC) and in the tail ganglia (TG). A set of 30 HCR probes (Molecular Instruments) and amplifiers labelled with Alexa 647 were used to detect the expression pattern of *G. pallida mod-1*. Confocal images are shown as maximum fluorescence projection images. Red fluorescence (**A, C**) and the superimposed image on a brightfield (**B, D**) are shown.

### Quipazine as a benchmark for chemicals that modulate MOD-1

Before commencing screening using the PhaGeM4 platform we sought to define a modulator of MOD-1 that could be used to benchmark subsequent “hits” from Pathogen Box. The developmental timing of *C. elegans* transgenics expressing either *C. elegans* or *G. pallida mod-1*, and co-expressing *gfp* in body wall muscle, was significantly delayed by quipazine as indicated by the reduction in fluorescence intensity which acts as a high throughput proxy forworm development [8] (Figure 2A). To validate quipazine as a MOD-1 modulator and lend confidence to the ability of this screening approach to yield MOD-1 “hits” we expressed *C. elegans* and *G. pallida* MOD-1 receptors in *X. laevis* oocytes for direct pharmacological analysis (Figure 2B). *C. elegans* MOD-1 was activated by 5-HT and the concentration-response relationship resulted in an EC_50_ value of 0.0929 µM. This EC_50_ is lower than that previously reported [7] however we also noted that the response exhibited very rapid desensitization. When expressed in Xenopus oocytes, *G. pallida* MOD-1 cRNAs reconstituted functional 5-HT receptors that were characterised by an EC_50_ value of 1.67 µM consistent with 5-HT acting as a cognate agonist to MOD-1. Quipazine also activated both *C. elegans* and *G. pallida* MOD-1 with EC_50_ values in a similar range of 10.28 µM and 31.57 µM, respectively. The maximum current elicited by quipazine was lower than the maximum elicited by 5-HT. Altogether, these results indicated that *C. elegans* and *G. pallida* MOD-1 form functional homomeric ligand-gated ion channel activated by 5-HT and at which quipazine acts as a partial agonist. This is consistent with the conclusion that quipazine delays development in the PhaGeM4 assay by acting as a modulator of MOD-1 expressed in the essential *C. elegans* pharyngeal neurone M4.

**Figure 2.**
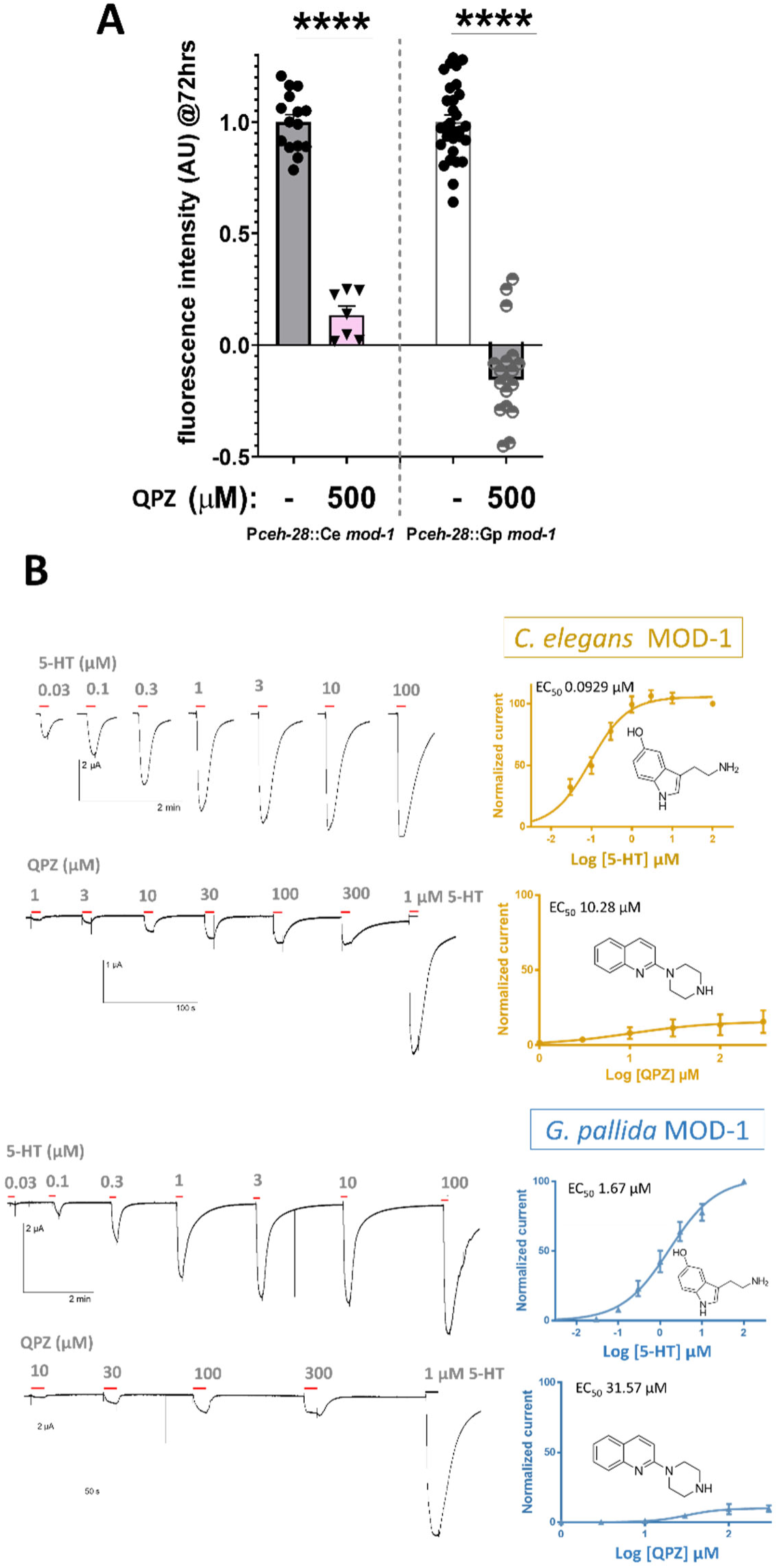
Quipazine as a benchmark for chemicals that modulate MOD-1. **A.** PhaGeM4_*mod-1* recombinant system shows quipazine (QPZ) as a potent partial agonist on both *C. elegans* and *G. pallida* MOD-1 receptor versions. The effect on development of 25 and 50 µM quipazine on strains expressing either P*ceh-28*::Ce *mod-1* or P*ceh-28*::Gp *mod-1* is indicated by the the fluorescence intensity of worms co-expressing *myo-3::gfp* as a readout of developmental progression. One way ANOVA (N=6-10 independent experiments) (**** p≤0.0001). **B.** Functional expression of both *C. elegans* and *G. pallida* MOD-1 versions in *X. laevis* oocytes. Representative current traces from an oocyte injected with cRNA of either Ce-MOD-1 (yellow) or Gp-MOD-1 (blue) challenged with the concentrations indicated of 5-HT and quipazine (QPZ). EC_50_ curves are shown to the right of the traces. N=7 oocytes for Ce-MOD-1 (5-HT), N=8 oocytes for Gp-MOD-1 (5-HT), N=5 oocytes for Ce-MOD-1 (QPZ), N=11 oocytes for Gp-MOD-1 (QPZ). **C.** Quipazine inhibits *C. elegans* motility in a *mod-1* dependent manner and more potently than 5-HT. One day old *C. elegans* were exposed to 2 mM 5-HT for 4 and 6 hrs, or 500 µM quipazine for 6 hours with and without 2 mM 5-HT. Each data point represents an independent assay of 10 worms. The right-hand graph shows that the effect of quipazine, and the greater inhibition seen with quipazine plus 5-HT, is abolished in the *mod-1* mutant background. One way ANOVA (N=4-10 independent experiments; ** p<0.01, **** p<0.0001 with respect to M9 control).

### Screening the Pathogen Box library in the PhaGeM4 *C. elegans* developmental assay identified several potential *C. elegans* MOD-1 “hits”

Having confirmed that the PhaGeM4 experimental platform identifies quipazine as a *boda fide* modulator of MOD-1, we proceeded to use *C. elegans* PhAGeM4 to screen for other MOD-1 modulators. In this screen we used a transgenic line stably expressing *C. elegans mod-1* in pharyngeal neurone M4. To conduct this screen, we ran control ‘non-drug’ and quipazine (QPZ) treatment experimental groups in parallel and used these as our benchmark for either no effect or potential MOD-1 modulation by the Pathogen Box compounds. Quipazine (25 µM, QPZ) reduced the control fluorescence by ∼75 % when recordings were made 72 hrs after allowing the inoculated eggs to develop in presence or absence of drug (Figure 3A). In a similar way we recorded the fluorescence of worms exposed to 25 µM of each of the Pathogen Box 400 compounds. We analysed the fluorescence from three independent experiments and calculated fluorescence intensity relative to the control for each drug. The complete set of data and relative fluorescence is shown (Figure 3A). These accumulated data are shown as a volcano plot (Figure 3B). Notably, we identified compounds that significantly inhibited growth in the transgenic worms, but which had no significant effect on N2 (wt). We prioritised 10 compounds with a significance value of p≤0.001 for further analysis as candidate modulators of MOD-1 (Figure 3B). In parallel to the fluorescence tracking in the PhaGeM4 bioassay, we monitored the OD_600 nm_ of *E. coli* OP50 added as a part of the monoxenic culture to assess the potential bacteriolytic effect exerted for individual compounds. This was an important control as a bacteriolytic effect may have an indirect effect on *C. elegans* growth by reducing food availability. We found a total of 7 compounds (Figure 3B) that induced an impairment in OP50 growth and that could be coupled to a negative effect in worm development. However, none of the 10 compounds that we selected for further analysis as potential modulators of MOD-1 had a bacteriolytic effect. In addition, we found 11 compounds (Figure 3B) where we observed an enhancement in fluorescence (p<0.05) compared to non-drug treatment. This increase in fluorescence after 72 hrs could be associated with a growth promoting effect, but no further analysis was made on this set of compounds.

**Figure 3.**
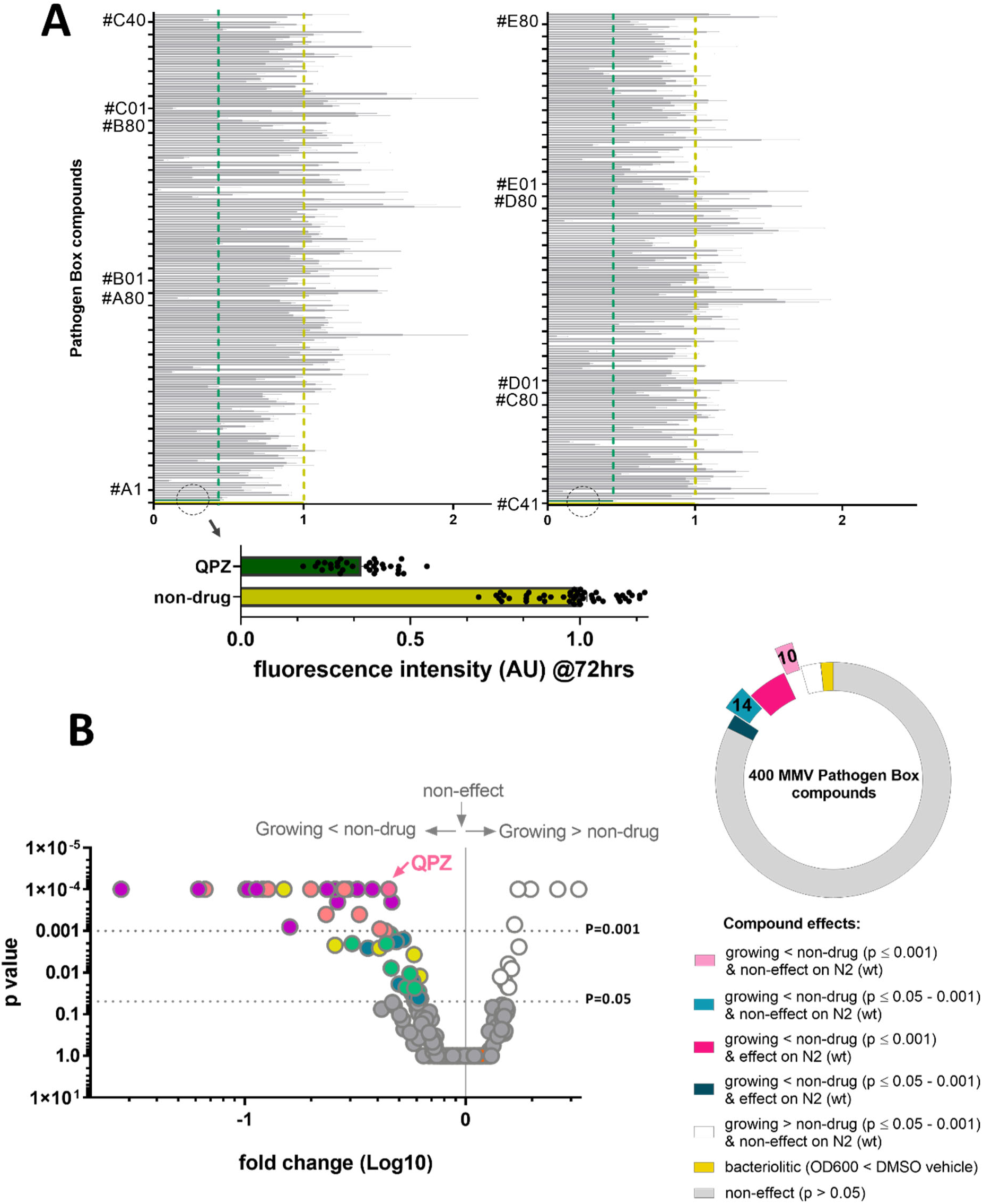
Screening of Pathogen Box compounds using the PhaGeM4 platform. **A**. Outcomes of the effects of the 400 individual compounds (25 µM) on development after 72 hrs from egg in the transgenic strain P*ceh-28*::*mod-1.* The level of fluorescence (AU) was measured as a readout of the growth in the presence of each compound. The dashed circle indicates the values for control ‘non-drug’ (yellow) and the benchmark quipazine (QPZ: green) and the vertical dashed lines in yellow and green indicate the benchmarks i.e., indicating the range for ‘no effect’ or potential MOD-1 modulation, respectively. Dashed circles represent a zoomed view of the base lines used as references (non-drug conditions and quipazine (QPZ) effect). Data are shown as mean ± s.e.m of 3 independent experiments for fluorescence intensity (AU) measured at 485nm/528nm (excitation/emission) at 72 hrs and normalised to non-drug. Statistical significance was determined using one way ANOVA. **B**. Volcano plot to highlight compounds with a significant effect on development (p=0.05-0.001) compared to the non-drug control. QPZ effect is indicated as reference; pale pink dots represent those compounds with an effect reducing growth compared to non-drug treatment (p≤0.001) plus non-effect on N2 (wt) strain that were identified as ‘hits’ for follow up; dark pink dots represent those compounds with an effect reducing growth compared to non-drug treatment (p≤0.001) that also had an effect on N2 (wt) strain and were not investigated further; green dots represent those compounds with an effect reducing growth compared to non-drug treatment (p=0.05-0.001) with (pale green) or without (dark green) an effect on N2 (wt) strain; yellow dots are compounds that had a bacteriolytic effect; white dots represent those compounds that produced an enhancement in fluorescence (P<0.05) compared to non-drug treatment, and could be associated with growth promoting. The circular diagram shows a classification of compounds with different level of effects on development in comparison to non-drug treatment, as well as compounds that were identified during the screening with bacteriolytic properties.

These preliminary results highlighted compounds within the Pathogen Box collection with potential effects selectively through the MOD-1 receptor (Figure 4A). This prompted us to perform re-testing using distinct batches of the 10 “hits” compounds by dissolving solid sample in DMSO from the selected.

**Figure 4.**
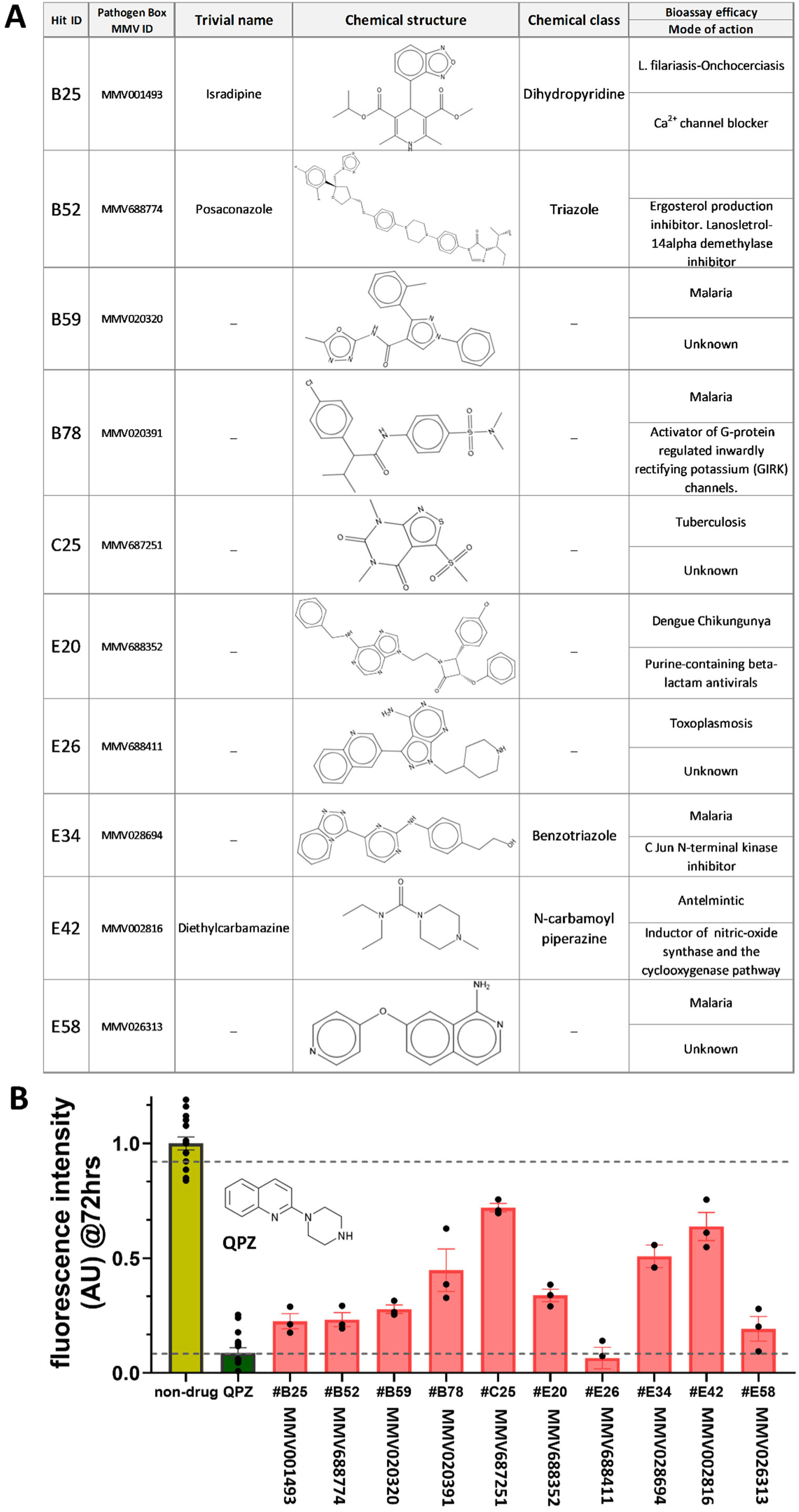
Selection and re-testing of hits with a significant MOD-1 dependent effect on developmental arrest. **A.** A list of the 10 selected hits out of the 400 compounds within the Pathogen Box library showing chemical information, data relating to their efficacy against parasites/viruses and information on mode of action. **B**. Re-testing of the 10 selected hits in the PhAGeM4 developmental assay using original MMV Pathogen Box dissolved stocks to confirm the effects on developmental arrest. All the ‘hits’ were tested at 25 µM with a final percentage of DMSO of 0.25%. The effect on individual hits is shown in comparison to non-drug (vehicle) and quipazine (QPZ, 25 µM) used as a reference compound. Data are shown as mean ± s.e.m of fluorescence intensity (AU) measured at 485nm/528nm (excitation/emission) at 72 hrs. N=3 independent experiments.

### Description of the *C. elegans* MOD-1 ‘hits’

Among the set of 10 ‘hits’, are compounds that are part of well-defined chemical classes (dihydropyridine, triazole, benzotriazole, N-carbamoyl piperazine), where the majority have established mode of actions. In contrast, there are other compounds, that although their efficacy in bioassays against various pathogens has been demonstrated, their mode of action remains unknown (B59-MMV020320, C25-MMV687251, E26-MMV688411, E58-MMV026313) (Figure 4A).

Upon re-testing all 10 ‘hits’ reproduced their efficacy in inhibiting *C. elegans* growth selectively in the transgenic expressing *mod-1* in M4 (Figure 4B).

### MOD-1 “hits” impair *G. pallida* motility

We investigated the efficacy of the ‘hits’ against *G. pallida* using motility assays with freshly hatched juveniles (J2 stage) (Figure 5A). We exposed J2s in microplates to 50 µM of each compound either for 24 hrs and scored motility. First, we tested 5-HT and quipazine both of which markedly reduced motility (Figure 5B) and then found that there was a significant reduction in motility for all of the ‘hit’ compounds (Figure 5C). These results suggest that the selected ‘hits’ exert an effective impairment on *G. pallida* motility and potentially impact its ability to infect a host plant.

**Figure 5.**
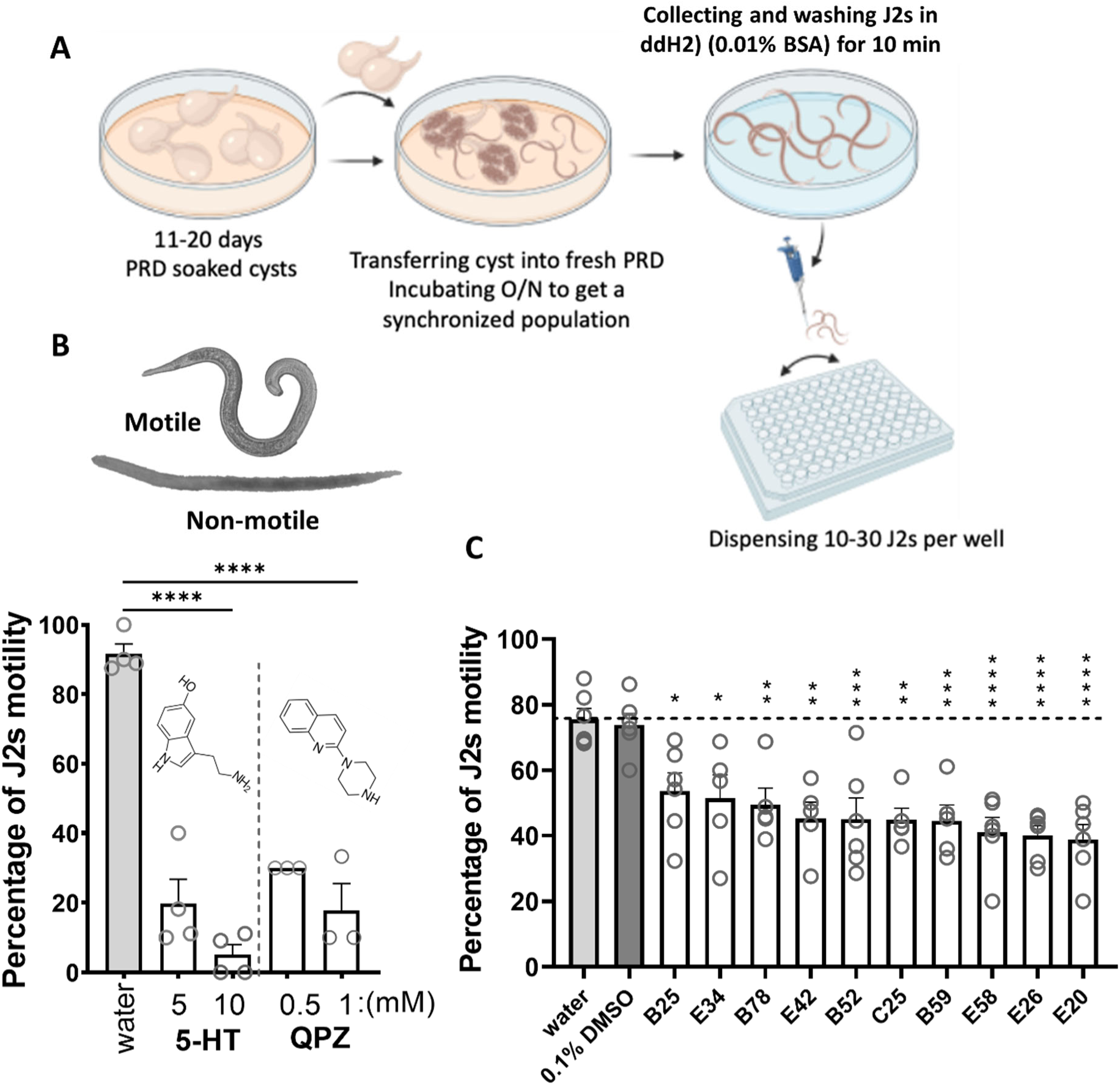
Testing the effectiveness of ‘hits’ in *G. pallida* using a motility bioassay. **A**. ∼20 *G. pallida* cysts were soaked for 11-20 days in potato root diffusate (PRD) in a 30 mm Petri dish. After 11-20 days when the J2 juveniles start emerging, the cysts were transferred into a new fresh dish with fresh (PRD), and the J2s emerged overnight were collected and washed with double distilled water before being dispensed into wells in a microplate containing 50 µM of each hit. Percentage of mobile juveniles was quantified after 4 hrs and 24 hrs. **B**. Quantification of *G. pallida* J2 motility after 24 hrs in chronic exposure with 5-HT and QPZ at the indicated doses. Representative images of the motile and non-motile juveniles are shown. Data are shown as mean ± s.e.m of percentage of mobile J2 with each data point representing an independent experiment. **C**. Quantification of *G. pallida* J2 motility after 24 hrs chronic exposure to selected Pathogen Box ‘hits’ at 50 µM. Two-way ANOVA with Bonferroni’s multiple comparisons (* p≤0.05, ** p≤0.01, *** p≤0.001, **** p≤0.0001).

### A subset of MOD-1 “hits” impairs *C. elegans* motility in a *mod-1* dependent manner

The screening described above allowed us to select 10 of the 400 Pathogen Box compounds for further investigation of their nematostatic or nematicidal potential. For *C. elegans* this was based on the understanding that 5-HT inhibits motility in a *mod-1* dependent manner [7]. Therefore, we tested the efficacy of the ‘hits’ in a *C. elegans* assay of motility in liquid, thrashing, and compared the effect on wild-type and *mod-1 (ok103),* the latter carrying a loss of function mutation in MOD-1. All the ‘hits’ except MMV026313 (E34) and MMV001493 (B25) reduced motility in a similar manner to the benchmark quipazine with the most effective being E42 (Figure 6). However, unlike the other compounds which had no effect on *mod-1(ok103)* supporting the contention that they act through MOD-1, E42 significantly reduced motility in both wild-type and *mod-1(ok103)* indicating that it also acts via another target or targets (Figure 6).

**Figure 6.**
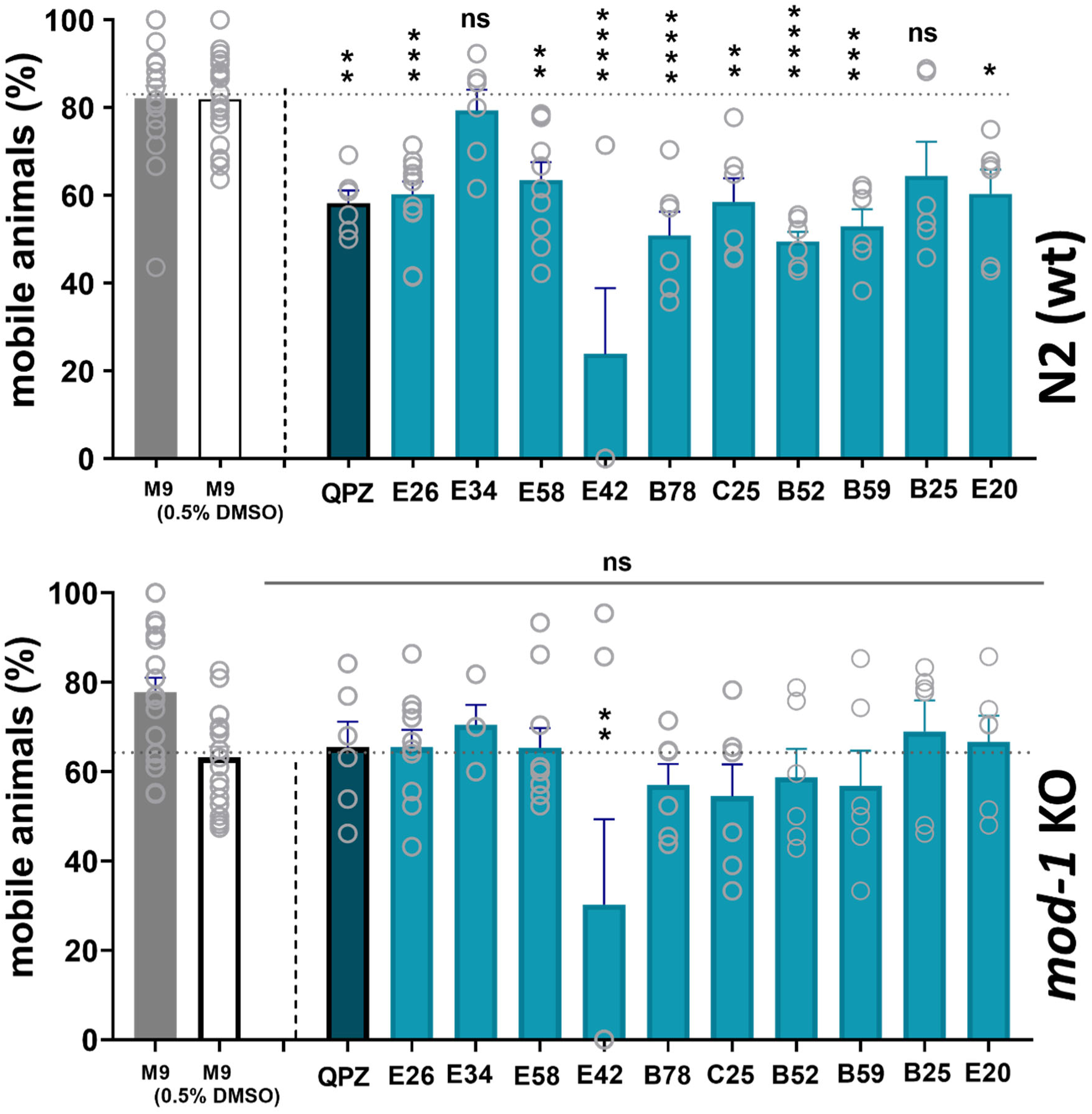
Testing the MOD-1 selective effect of ‘hits’ using a *C. elegans* motility bioassay. **A**. 20 L4+1-day old N2 (wt) or **B**. *mod-1(ok103)* worms were incubated for 6 hrs in vehicle or 25 µM of the compound as indicated. The effect on thrashing was evaluated by quantification of the percentage of mobile worms after 6 hrs. Data are shown as mean ± s.e.m. Each data point represents an independent experiment. Two-way ANOVA with Bonferroni’s multiple comparisons (* p≤0.05, ** p≤0.01, *** p≤0.001, **** p≤0.0001).

### A subset of the *C. elegans* MOD-1 ‘hits’ impair slow development in PhaGeM4 expressing *G. pallida mod-1*

We next wanted to test whether the potential *C. elegans* MOD-1 modulators, i.e., the 10 ‘hits’ selected from the preliminary screen, had efficacy at *G. pallida* MOD-1. We therefore compared the effect of the set of 10 ‘hits’, at two concentrations (25 µM and 50 µM) in PhaGeM4, by incubating eggs from the transgenic line P*ceh-28*::*Gp mod-1* with each of the ten ‘hit’ compounds. For this second comparative re-testing we used compounds provided from the Pathogen Box partner Evotec Toulouse (France) as solid phase and then constituted in DMSO. MMV687251 (C25), MMV688774 (B52) and MMV002816 (E42) significantly inhibited development in an apparently concentration-dependent manner (Figure 7). C25 completely arrested growth at 25 and 50 µM therefore we tested it at lower concentrations of 1 and 10 µM which were both without effect indicating a threshold concentration of between 10 and 25 µM. All the other compounds tested, except MMV026313 (E34), and MMV688352 (E20), had some significant effect on development. However, this was not dose-dependent and therefore not convincing in terms of a genuine effect of MOD-1 modulation (Supplementary Figure 1).

**Figure 7.**
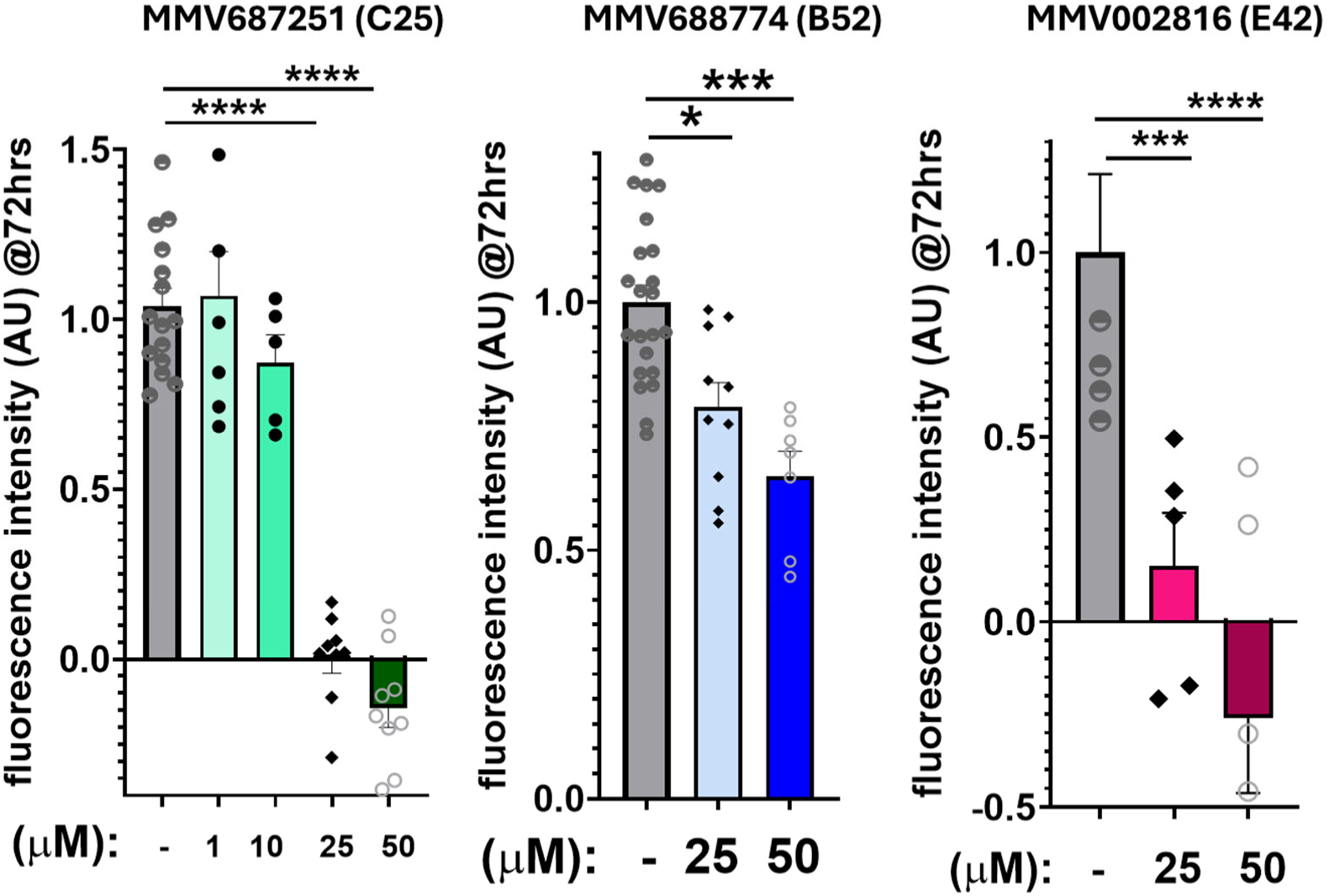
Testing hits on MOD-1 receptor from *G. pallida* in the *C. elegans* PhaGeM4 assay identifies three compounds with efficacy. The effect on development of each compound on the transgenic strain expressing *G. pallida mod*-1 i.e., carrying P*ceh-28*::Gp *mod-1.* Data are shown as mean ± s.e.m of normalised fluorescence intensity (AU) measured at 485nm/528nm (excitation/emission) at 72 hrs. Two-way ANOVA with Bonferroni’s multiple comparisons (* p≤0.05, ** p≤0.01, *** p≤0.001, **** p≤0.0001).

### Four of the MOD-1 “hits” impair the ability of *G. pallida* J2s to invade host potato roots

A nematode root invasion assay was employed to evaluate the impact of quipazine and the selected ‘hits’ on the ability of *G. pallida* J2 to invade host potato roots. The total number of *G. pallida* J2s able to infect host roots after 24 hrs exposure to individual ‘hits’ was quantified at 7 days post-infection (dpi). With the exception of quipazine, which dissolves in water, all the stock solutions were prepared in 100% DMSO. Therefore, we first tested the effect of 0.1%, 0.3% and 0.9% DMSO on the ability of *G. pallida* J2 to infect host potato roots. There was no significant difference in the ability to infect potato host roots between the control *G. pallida* J2s that were treated with water and *G. pallida* J2s treated with any of the DMSO concentrations tested (Supplementary Figure 2). We only employed solutions containing ≤ 0.9%DMSO in subsequent assays. As the compounds had solubilities ranging from 50-450 µM in DMSO, the maximum testable concentration for each ‘hit’ varied.

Quipazine reduced the ability of *G. pallida* J2 to infect host potato roots in a concentration-dependent manner (Figure 8A) and at 2mM this appeared to effect complete protection from invasion. At 450 µM, B52, C25, E20, E42 all significantly reduced the ability of *G. pallida* J2s to invade host potato roots (Figure 8B). In the case of E42, even concentrations as low as 16.7 µM impacted invasion (Supplementary Figure 3). A reduced trend could be observed from exposure of J2s to concentrations as low as 16.7 µM of B52 and 150 µM of C25 (Supplementary Figure 3). Exposure of J2s to B25, B59, B78, E26, E34 and E58 did not affect root invasion at any of the concentrations tested (Supplementary Figure 4). Due to solubility limits higher concentrations of these ‘hits’ could not be employed. However, we concluded that at least 40% of the MOD-1 ‘hits’ identified during the PhaGeM4 screen impact the ability of *G. pallida* J2s to invade host roots.

**Figure 8.**
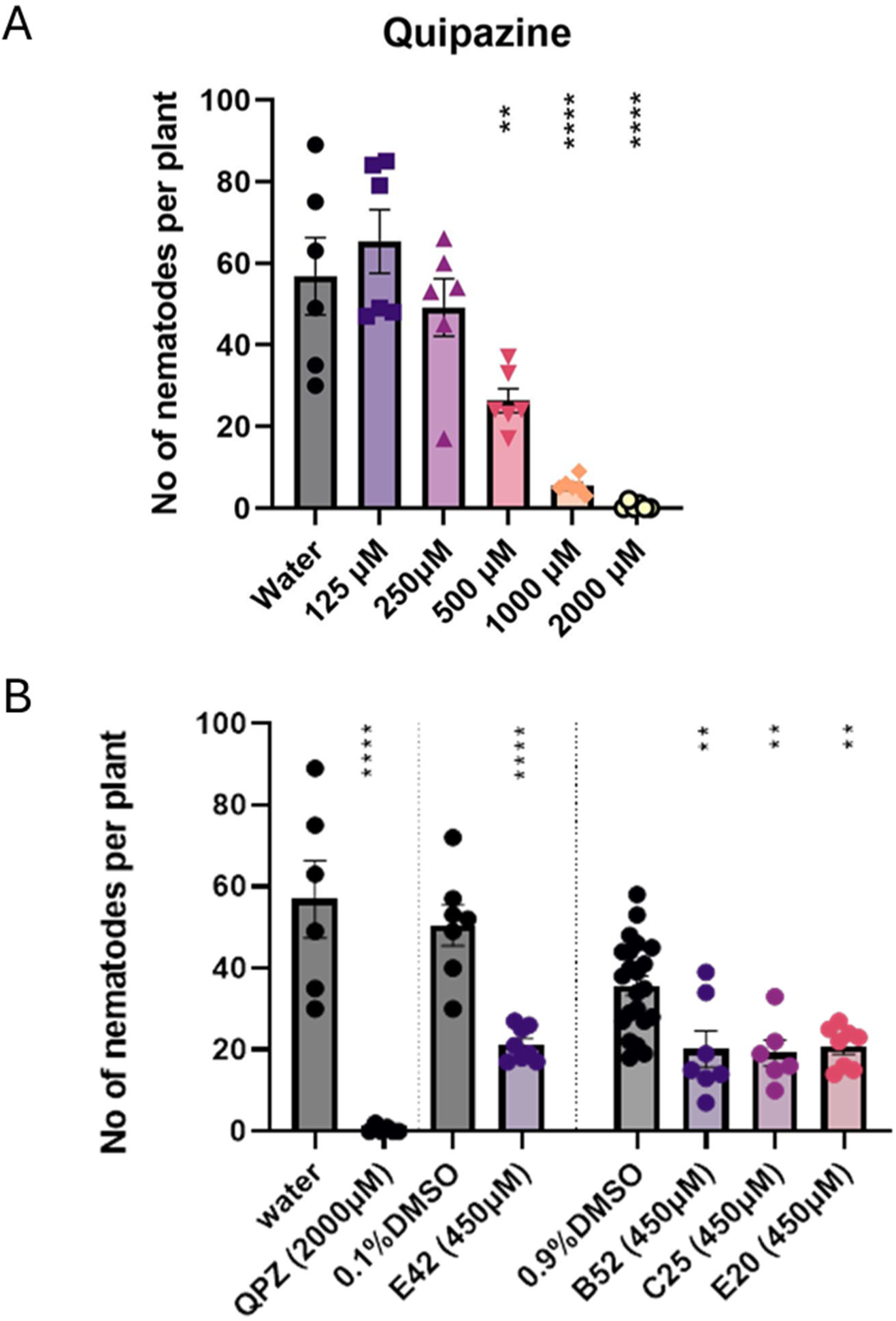
Exposure to quipazine, E42, B52, C25 and E20 reduces *G. pallida* J2s ability to invade host potato roots. Hatched *G. pallida* J2s were exposed for 24h prior to either test compound or water. Potato plants cv Desiree grown in pouches were inoculated with the J2s with 5 infection points/root system and ∼ 25 J2/ infection point. The nematode root invasion was measured as the total number of nematodes that were present in the roots 7 dpi as identified by staining with acid fuchsin. **A.** *G. pallida* J2 exposed to concentrations of quipazine ≥ 500µM showed a significant reduction in their ability to infect host potato roots. **B.** As for A, except *G. pallida* J2s were exposed to either 450 µM B52, C25, E20, or E42. Data are shown as mean ± standard error of the mean. One way ANOVA followed by Dunnett post hoc in case of B52, C25 and E20. Student’s t-test for E42 and quipazine (** p≤0.01, **** p≤0.0001) (n=6-8).

## DISCUSSION

Here we have provided the first characterisation of the 5-HT-gated chloride MOD-1 in the plant parasitic nematode *G. pallida* as a step towards exploring its potential as a novel nematicide target. Previously, we had identified *mod-1* from two potato cyst nematodes, *G. pallida* and *Globodera rostochiensis* which predicts receptors with 46.1 and 45.8% amino acid identity to *C. elegans* MOD-1, respectively [6]. Although highly homologous the predicted protein for *G. pallida* is longer than *C. elegans*, 656 compared to 489 amino acids, and exhibits interesting differences. These are largely reflected in the expanded third cytoplasmic loop, a domain associated with trafficking, scaffolding and signalling cross -talk of the superfamily of cys-loop receptors [20] of which MOD1 is a member [21]. In addition, there is a clear extension in the extracellular C-terminal extensions that acts as an important determinant of activation in this class of receptor. This suggests there may be some variation in the otherwise conserved determinant of the receptors’ function [22]. These differences may reflect the variation in functional pharmacology of the *C. elegans* and *G. pallida* recombinantly expressed *mod-1* analysed in *Xenopus* oocytes. Interestingly not all the *C. elegans* hits were confirmed in *G. pallida* suggesting there may be differences in pharmacology between the two species. Nonetheless, like *C. elegans* MOD-1, it is activated by 5-HT and quipazine acts as a partial agonist. Moreover, the expression pattern in *G. pallida* appears similar to that in *C. elegans* consistent with a conserved functional role. Notably, *G. pallida mod-1* is expressed in the ventral nerve cord which may reflect a key role in regulating motility as described in *C. elegans* [7] and lending confidence to the rationale of exploring it as a nematicide target.

We used the 5-HT receptor agonist quipazine as the benchmark for screening purpose having previously shown that it is an effective agonist for *C. elegans* MOD-1 [8]. Quipazine phenocopies the *mod-1* dependent inhibitory effect of 5-HT on *C. elegans* motility reinforcing the rationale that targeting *mod-1* may have a paralytic action. Quipazine also markedly inhibited *C. elegans* development in transgenics expressing *G. pallida mod-1* in M4, consistent with its observed agonist action at this receptor. In this regard, the clear and striking efficacy of quipazine in eliciting a concentration-dependent inhibition of J2 root invasion strongly aligns with the contention that targeting MOD-1 in this way has potential for crop protection. Indeed, it suggests that quipazine-like compounds may be worth further investigation in this regard.

Using quipazine as our benchmark we screened the Pathogen Box set of compounds for efficacy against *C. elegans* MOD-1 in PhaGeM4. This yielded 10 ‘hits’ all of which affected *C. elegans* motility in *mod-1* dependent manner except for E42, MMV002816. This compound is the antifilarial compound, diethylcarbamzine. It had a significant effect in all the bioassays however it also inhibited motility in the *C. elegans* mutant *mod-1* (*ok103*) suggesting that it exerts its action on motility through a mechanism other than, or in addition to, a modulation of MOD-1. The antifilarial mode of action for diethethlycarbamzine is not entirely clear. It has been suggested that it may interact with a TRP channel in both the nematode and the host [23]. It is interesting to speculate that an interaction with MOD-1 may also contribute to its anthelmintic action. In terms of plant parasitic nematode control, diethylcarbamazine was very effective at preventing *G. pallida* J2 root invasion and is worthy of follow up.

From the screen and associated bioassays, two of the original *C. elegans* hits met all the criteria for MOD-1 dependent inhibition of root invasion (Table 1). C25, MMV687251 is a vancomycin-like compound [24] that has been identified as having potential in biofilm control. To our knowledge, it has not previously been identified in screens for nematicides or anthelmintics. B52, MMV688774, is the antifungal agent posaconazole [25]. As an antifungal, its mode of action is an inhibitor of ergosterol synthesis. Our data suggest that this chemical moiety also interacts with MOD-1 to impair the viability of plant parasitic nematodes and this merits further investigation.

**Table 1.** Summary of the progression of ‘hit’ compounds through the MOD-1 screening pipeline. The top row names the assays that were carried out for each of the hits, starting with the *C. elegans* PhaGeM4 assay in which *C. elegans mod-1* was expressed in M4, then the *G. pallida* motility and root invasion assay, followed by the *C. elegans* motility assay and a column indicating whether the inhibitory effect of the compound on *C. elegans* motility was dependent on *mod-1* (i.e., not observed in the *mod-1* mutant *ok103*). The last two column on the right represents results from the *G. pallida* PhaGeM4 assay in which *G. pallida mod-1* was expressed in *C. elegans* M4. The results are colour coded; green indicates an effect was observed and orange indicates either no effect or that the effect was not dose-dependent and therefore unlikely to be a genuine effect of the compound. QUIP is the benchmark quipazine. The table shows that the *G. pallida* PhaGeM4 assay predicts 4 compounds (including quipazine) that have an inhibitory effect on J2 root invasion of which one, E42, is also acting on a target other than MOD-1.

| Identifier | MMV code | *C. elegans* PhaGeM4 | *G. pallida* motility | *C. elegans* motility | *mod-1* dependent? | *G. Pallida* PhaGeM4 | Root invasion |
| --- | --- | --- | --- | --- | --- | --- | --- |
| QUIP |  |  |  |  |  |  |  |
| C25 | MMV687251 |  |  |  |  |  |  |
| B52 | MMV688774 |  |  |  |  |  |  |
| E42 | MMV002816 |  |  |  | no |  |  |
| E20 | MMV688352 |  |  |  |  | no effect |  |
| B25 | MMV001493 |  |  | no effect |  | not dose dependent | no effect |
| B78 | MMV020391 |  |  |  |  | not dose dependent | no effect |
| E58 | MMV026313 |  |  |  |  | not dose dependent | no effect |
| B59 | MMV020320 |  |  |  |  | not dose dependent | no effect |
| E26 | MMV688411 |  |  |  |  | not dose dependent | no effect |
| E34 | MMV028694 |  |  | no effect |  | no effect | no effect |

Overall, the pipeline we developed to assess Pathogen Box compounds for efficacy against plant parasitic nematodes lends confidence that transgenically expressing MOD-1 in *C. elegans* pharyngeal neurone M4 can deliver new chemical leads. More generally, this proof-of-concept study provides a new tool to screen compound libraries for lead chemicals of interest. Indeed, this approach could be developed for other ligand-gated ion channels that are candidate nematicide or anthelmintic targets providing a facile whole organism screen for novel chemistry.

## Funding

F.C. and K.M. were supported by Biotechnology and Biological Sciences (BBSRC) grant number BB/T002867/1.

## Acknowledgements

Some *C. elegans* strains were provided by the CGC, which is funded by NIH Office of Research Infrastructure Programs (P40 OD010440). We are grateful to Medicines for Malaria Venture (MMV; https://www.mmv.org/mmv-open/pathogen-box/about-pathogen-box) for The Pathogen Box compounds and to Evotec Toulouse (France) Evotec for supplying lyophilized samples.

## Competing interests

Dr. Claude L. Charvet is an employee of MSD Animal Health Innovation GmbH. All other authors declare no conflict of interest.

## REFERENCES

[1] A. Taylor, Nematocides and nematicides-a history, Nematropica, (2003) 225–232.

[2] A.S.S. Schleker, M. Rist, C. Matera, A. Damijonaitis, U. Collienne, K. Matsuoka, S.S. Habash, K. Twelker, O. Gutbrod, C. Saalwächter, M. Windau, S. Matthiesen, T. Stefanovska, M. Scharwey, M.T. Marx, S. Geibel, F.M.W. Grundler, Mode of action of fluopyram in plant-parasitic nematodes, Scientific Reports, 12 (2022) 11954.

[3] G. Lahm, J. Desaeger, B. Smith, T. Pahutski, M. Rivera, T. Meloro, R. Kucharczyk, R. Lett, A. Daly, B. Smith, D. Cordova, T. Thoden, J. Wiles, The discovery of fluazaindolizine: A new product for the control of plant parasitic nematodes, Bioorg Med Chem Lett, 27 (2017).

[4] Y. Oka, S. Shuker, N. Tkachi, Nematicidal efficacy of MCW-2, a new nematicide of the fluoroalkenyl group, against the root-knot nematode Meloidogyne javanica, Pest Management Science: formerly Pesticide Science, 65 (2009) 1082–1089.

[5] J. Kearn, E. Ludlow, J. Dillon, V. O’Connor, L. Holden-Dye, Fluensulfone is a nematicide with a mode of action distinct from anticholinesterases and macrocyclic lactones, Pestic Biochem Physiol, 109 (2014) 44–57.

[6] A. Crisford, F. Calahorro, E. Ludlow, J.M.C. Marvin, J.K. Hibbard, C.J. Lilley, J. Kearn, F. Keefe, P. Johnson, R. Harmer, P.E. Urwin, V. O’Connor, L. Holden-Dye, Identification and characterisation of serotonin signalling in the potato cyst nematode Globodera pallida reveals new targets for crop protection, Plos Pathog, 16 (2020).

[7] R. Ranganathan, S.C. Cannon, H.R. Horvitz, MOD-1 is a serotonin-gated chloride channel that modulates locomotory behaviour in C. elegans, Nature, 408 (2000) 470–475.

[8] F. Calahorro, M. Chapman, K. Dudkiewicz, L. Holden-Dye, V. O’Connor, PharmacoGenetic targeting of a C. elegans essential neuron provides an in vivo screening for novel modulators of nematode ion channel function, Pesticide Biochemistry and Physiology, 186 (2022) 105152–105152.

[9] N. Rodriguez Araujo, G. Hernando, J. Corradi, C. Bouzat, The nematode serotonin-gated chloride channel MOD-1: A novel target for anthelmintic therapy, J Biol Chem, (2022) 102356.

[10] L. Avery, H.R. Horvitz, A Cell That Dies during Wild-Type C-Elegans Development Can Function as a Neuron in a Ced-3 Mutant, Cell, 51 (1987) 1071–1078.

[11] C.G.L. Veale, Unpacking the Pathogen Box-An Open Source Tool for Fighting Neglected Tropical Disease, Chemmedchem, 14 (2019) 386–453.

[12] S. Duffy, M.L. Sykes, A.J. Jones, T.B. Shelper, M. Simpson, R. Lang, S.A. Poulsen, B.E. Sleebs, V.M. Avery, Screening the Medicines for Malaria Venture Pathogen Box across Multiple Pathogens Reclassifies Starting Points for Open-Source Drug Discovery, Antimicrob Agents Ch, 61 (2017).

[13] P. Thorpe, S. Mantelin, P.J. Cock, V.C. Blok, M.C. Coke, S. Eves-van den Akker, E. Guzeeva, C.J. Lilley, G. Smant, A.J. Reid, K.M. Wright, P.E. Urwin, J.T. Jones, Genomic characterisation of the effector complement of the potato cyst nematode Globodera pallida, BMC Genomics, 15 (2014) 923.

[14] A.L. Sperling, S. Eves-van den Akker, Whole mount multiplexed visualization of DNA, mRNA, and protein in plant-parasitic nematodes, Plant Methods, 19 (2023) 139.

[15] H.M. Choi, C.R. Calvert, N. Husain, D. Huss, J.C. Barsi, B.E. Deverman, R.C. Hunter, M. Kato, S.M. Lee, A.C. Abelin, A.Z. Rosenthal, O.S. Akbari, Y. Li, B.A. Hay, P.W. Sternberg, P.H. Patterson, E.H. Davidson, S.K. Mazmanian, D.A. Prober, M. van de Rijn, J.R. Leadbetter, D.K. Newman, C. Readhead, M.E. Bronner, B. Wold, R. Lansford, T. Sauka-Spengler, S.E. Fraser, N.A. Pierce, Mapping a multiplexed zoo of mRNA expression, Development, 143 (2016) 3632–3637.

[16] S. Brenner, Genetics of Caenorhabditis-Elegans, Genetics, 77 (1974) 71–94.

[17] P.E. Urwin, C.J. Lilley, M.J. McPherson, H.J. Atkinson, Resistance to both cyst and root-knot nematodes conferred by transgenic Arabidopsis expressing a modified plant cystatin, Plant J, 12 (1997) 455–461.

[18] A. Blanchard, F. Guegnard, C.L. Charvet, A. Crisford, E. Courtot, C. Sauve, A. Harmache, T. Duguet, V. O’Connor, P. Castagnone-Sereno, B. Reaves, A.J. Wolstenholme, R.N. Beech, L. Holden-Dye, C. Neveu, Deciphering the molecular determinants of cholinergic anthelmintic sensitivity in nematodes: When novel functional validation approaches highlight major differences between the model Caenorhabditis elegans and parasitic species, Plos Pathog, 14 (2018).

[19] G. Gürel, M.A. Gustafson, J.S. Pepper, H.R. Horvitz, M.R. Koelle, Receptors and other signaling proteins required for serotonin control of locomotion in *Caenorhabditis elegans*, Genetics, 192 (2012) 1359–1371.

[20] C.M. Noviello, A. Gharpure, N. Mukhtasimova, R. Cabuco, L. Baxter, D. Borek, S.M. Sine, R.E. Hibbs, Structure and gating mechanism of the α7 nicotinic acetylcholine receptor, Cell, 184 (2021) 2121–2134.e2113.

[21] A.K. Jones, D.B. Sattelle, The cys-loop ligand-gated ion channel gene superfamily of the nematode, Caenorhabditis elegans, Invert Neurosci, 8 (2008) 41–47.

[22] S.A. Pless, L.G. Sivilotti, A tale of ligands big and small: an update on how pentameric ligand-gated ion channels interact with agonists and proteins, Curr Opin Physiol, 2 (2019) 19–26.

[23] P.D.E. Williams, S.S. Kashyap, A.P. Robertson, R.J. Martin, Diethylcarbamazine elicits calcium signals by activation of Brugia malayi TRP-2b channels heterologously expressed in HEK293 cells, Sci Rep, (2025).

[24] V. Bhandari, S. Chakraborty, U. Brahma, P. Sharma, Identification of anti-staphylococcal and anti-biofilm compounds by repurposing the Medicines for Malaria Venture Pathogen Box, Front Cell Infect Microbiol, 8 (2018) 365.

[25] L. Heimark, P. Shipkova, J. Greene, H. Munayyer, T. Yarosh-Tomaine, B. DiDomenico, R. Hare, B.N. Pramanik, Mechanism of azole antifungal activity as determined by liquid chromatographic/mass spectrometric monitoring of ergosterol biosynthesis, J Mass Spectrom, 37 (2002) 265–269.

