## Supplemental figures for "Potential nematicides identified through targeted *in vivo* screening of a 5-HT-gated chloride channel, MOD-1"

**Supplementary Figure 1. Testing hits on MOD-1 receptor from *G. pallida* in the *C. elegans* PhaGeM4 assay identifies compounds that are either without effect or for which the effect is not concentration-dependent.** The effect on development of each compound on the transgenic strain expressing *G. pallida mod*-1 i.e., carrying P*ceh-28*::Gp *mod-1.* Data are shown as mean ± s.e.m of normalised fluorescence intensity (AU) measured at 485nm/528nm (excitation/emission) at 72 hrs. Two-way ANOVA with Bonferroni's multiple comparisons (* p≤0.05, ** p≤0.01, *** p≤0.001, **** p≤0.0001).

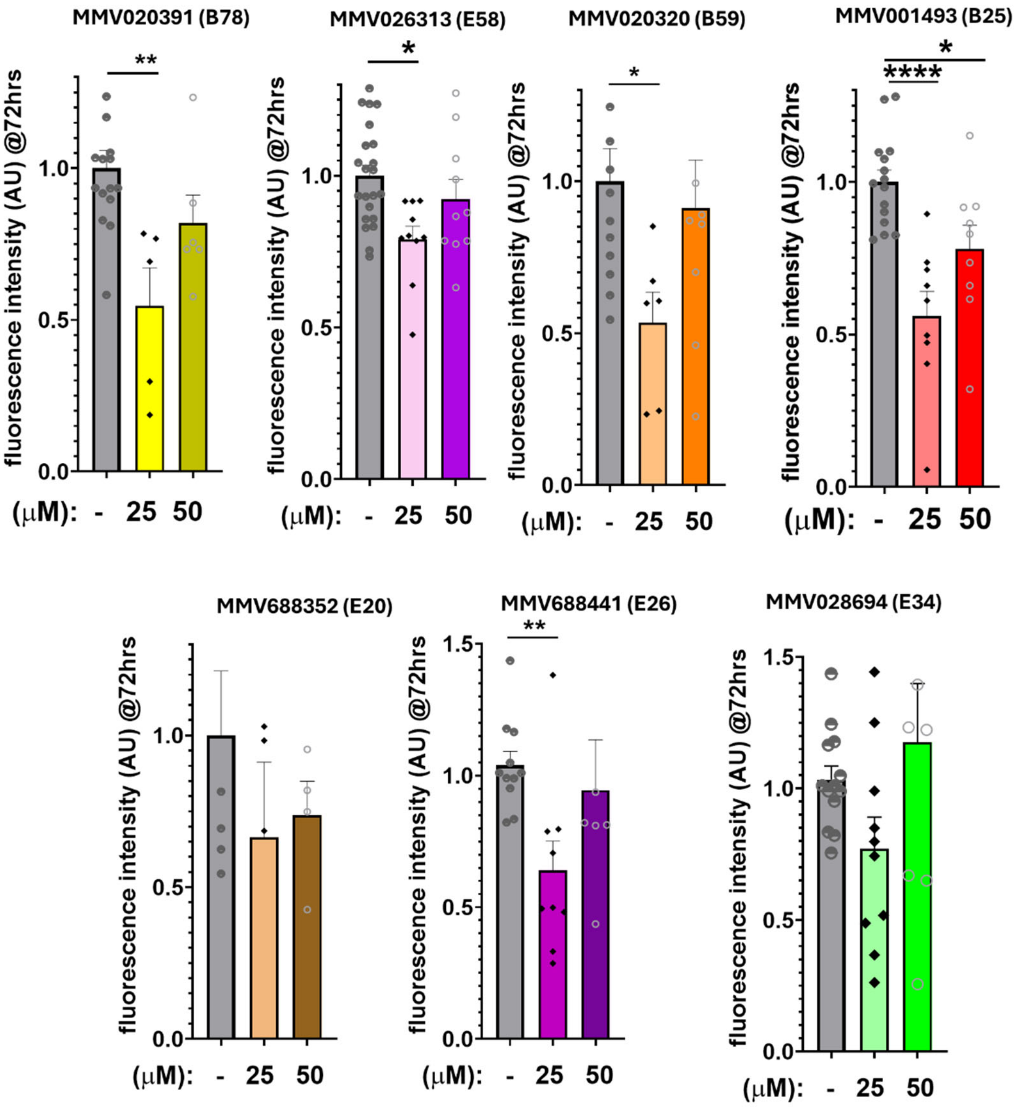

**Supplementary Figure 2. Exposure to 0.1% DMSO, 0.3% DMSO or 0.9% DMSO did not affect the ability of *G.pallida* J2s to invade host potato roots**. Hatched *G.pallida* J2s were exposed for 24h prior inoculation **0**.1% DMSO, 0.3% DMSO or 0.9% DMSO and water, in case of control J2s. Potato plants cv Desiree grown in pouches were inoculated with *G.pallida* J2s treated as described above, with 5 infection points/root system and ~ 25 J2/ infection point. The nematode invasion measured as total number of nematodes able to infect potato host roots was determined 7 dpi following staining with acid fuchsin. *G.pallida* J2 exposed to 0.1% DMSO, 0.3% DMSO or 0.9% DMSO did not significantly affect the ability of J2s to invade host potato roots when compared to *G.pallida* J2s exposed to water only. Data is shown as means, and error bars are SEM. One way ANOVA followed by Dunnett post hoc (n=6-8).

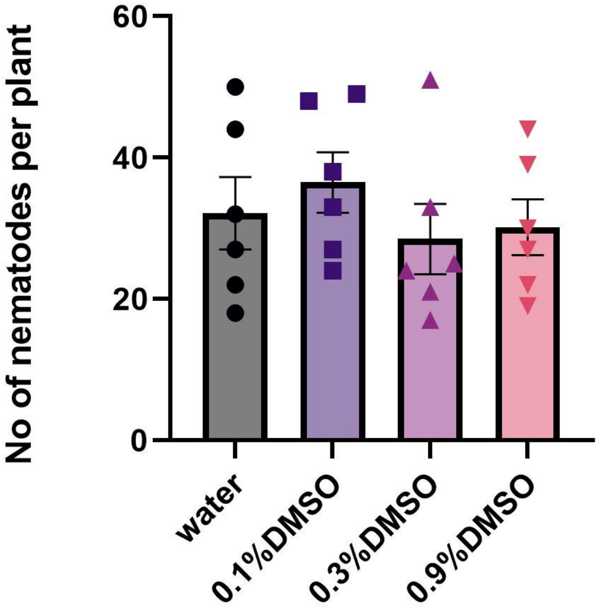

**Supplementary Figure 3. A concentration range was tested for B52, C25, E20 and E42.** Hatched *G. pallida* J2s were exposed for 24h prior to inoculation to different concentrations of B52, C25, E20 and E42 or 0.9% DMSO, 0.1% DMSO and water in case of control J2s. Potato plants cv Desiree grown in pouches were inoculated with *G. pallida* J2s treated as described above, with 5 infection points/root system and ~ 25 J2/ infection point. The nematode invasion measured as total number of nematodes able to infect potato host roots was determined 7 dpi following staining with acid fuchsin**.** While only *G. pallida* J2 exposed to 450 µM B52 showed a significant reduction in their ability to infect host potato roots, a reduced trend can be observed from an exposure of J2s to concentrations of B52 ≥16.7µM**.** While only *G. pallida* J2 exposed to 1350 µM C25 showed a significant reduction in their ability to infect host potato roots a reduced trend can be observed from an exposure of J2s to concentrations of C25 ≥150µM**.** Only *G. pallida* J2 exposed to 450 µM E20 showed a significant reduction in their ability to infect host potato roots. *G. pallida* J2 exposed to concentrations of E42 ≥ 16.7 µM showed a significant reduction in their ability to infect host potato roots with a reduced trend being observed from an exposure of J2s to 16.7 µM E422**.** Data are shown as means, and error bars are SEM. One way ANOVA followed by Dunnett post hoc. (* p≤0.05, ** p≤0.01, *** p≤0.001,**** p≤0.0001) (n=6-8).

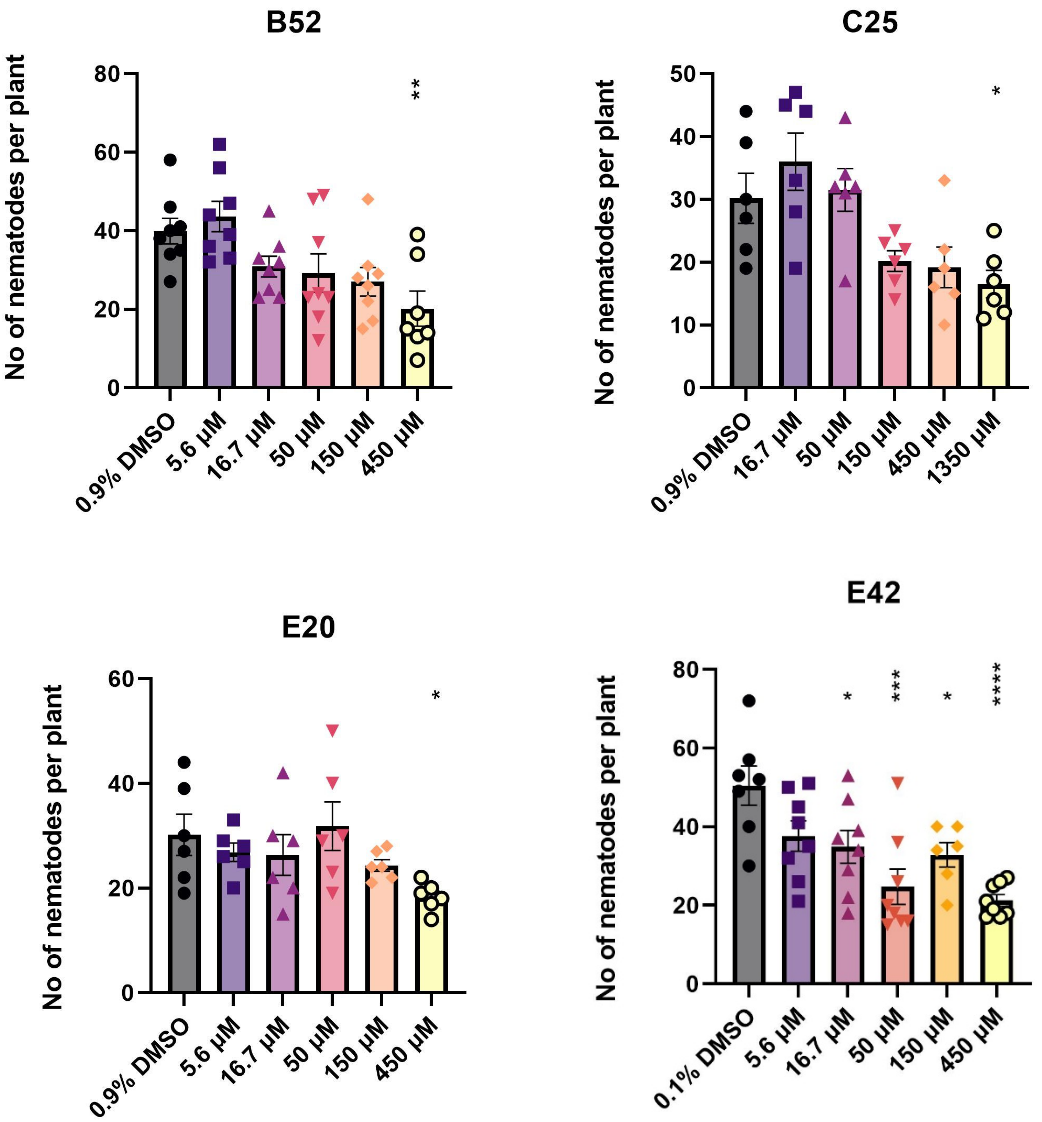
B52
C25
E20
E42
No of nematodes per plant
0.9% DMSO
0.1% DMSO
5.6 µM
16.7 µM
50 µM
150 µM
450 µM
1350 µM

**Supplementary Figure 4. Exposure to B25, B59, B78, E26, E34 and E58 did not affect the ability of *G. pallida* J2s to invade host potato roots**. Hatched *G . pallida* J2s were exposed for 24h prior inoculation to B25, B59, B78, E26, E34 or 0.9% DMSO, in case of control J2s. Potato plants cv Desiree grown in pouches were inoculated with *G. pallida* J2s treated as described above, with 5 infection points/root system and ~ 25 J2/ infection point. The nematode invasion measured as total number of nematodes able to infect potato host roots was determined 7 dpi following staining with acid fuchsin. *G. pallida* J2 exposed to B25, B59, B78, E26, E34 and E58 did not significantly affect the ability of J2s to invade host potato roots. Data are shown as means, and error bars are SEM. One way ANOVA followed by Dunnett post hoc (n=6-8).

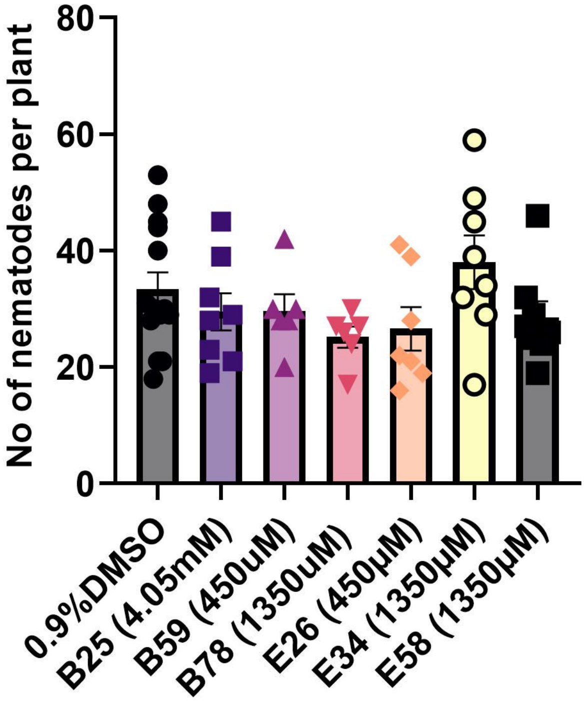
No of nematodes per plant
80
60
40
20
0
0.9%DMSO
B25 (4.05mM)
B59 (450uM)
B78 (1350uM)
E26 (450µM)
E34 (1350µM)
E58 (1350µM)

52
